# Identification of functional non-coding variants affecting Alzheimer’s disease risk by Massively Parallel Reporter Assay

**DOI:** 10.64898/2026.04.30.721912

**Authors:** Adam D. Hudgins, Hyun-Kyung Chang, Seung-Soo Kim, Jiping Yang, Di Guan, Xizhe Wang, Yizhou Zhu, Yousin Suh

## Abstract

The majority of genetic risk variants for late-onset Alzheimer’s disease (LOAD) reside within non-coding genomic regions, suggesting they exert pathogenic effects by disrupting transcriptional regulatory programs. To systematically identify functional variants, we performed an unbiased, high-throughput Massively Parallel Reporter Assay (MPRA) in the THP-1 human monocytic cell line, screening 2,231 SNPs across 24 LOAD GWAS loci. We identified 62 variants exhibiting significant allele-specific transcriptional regulatory output, including rs636317 in the *MS4A* locus. The risk allele of rs636317 disrupts a CTCF binding motif, altering fine-scale chromatin looping. To validate these findings, we employed CRISPR/Cas9 to generate an allele-specific deletion of the rs636317 T allele in H9 human embryonic stem cells. Monocytes differentiated from these edited cells displayed increased expression of *MS4A4E*, *MS4A6A*, and *TREM2*, along with highly elevated levels of soluble TREM2 (sTREM2). These results provide a mechanistic link between a non-coding genetic variant, transcriptional regulation of the *MS4A* family, and *TREM2*-mediated immune responses in Alzheimer’s disease pathogenesis.

## Introduction

Alzheimer’s Disease (AD) is a prevalent, age-related neurodegenerative disorder and the leading cause of dementia globally. While genome-wide association studies (GWAS) have successfully identified over 40 susceptibility loci for sporadic late-onset AD (LOAD), the vast majority of these implicated variants reside within non-coding genomic regions^1–6^. This localization significantly complicates efforts to identify the underlying causal genes. Furthermore, extensive genetic linkage disequilibrium often results in the co-inheritance of multiple significant variants within a single locus, making it difficult to discern which specific variant fundamentally drives disease susceptibility. Consequently, pinpointing true causal variants and characterizing their mechanistic contributions to LOAD pathogenesis remains a critical challenge in the post-genomic era.

Recent studies have presented evidence that non-coding GWAS variants can contribute to disease risk and pathogenesis by altering cell- and tissue-specific transcriptional regulatory programs^3,7^. To meet this challenge, an experimental method known as the Massively Parallel Reporter Assay (MPRA) has been developed as a high-throughput, unbiased approach to link non-coding variants to transcriptional outputs^8–11^. Through the use of MPRA, genomic regions harboring GWAS non-coding risk variants, or those in linkage disequilibrium (LD), are screened for transcriptional control functions.

Choosing the correct cellular model in which to carry out the MPRA is critical, as reporter assays display cell-type-specific activity^12^. Previous analyses have shown that non-coding LOAD GWAS variants are enriched in the transcriptional enhancers of myeloid lineage cells, including monocytes, macrophages, and microglia^13–15^. Research into the cellular mechanisms of LOAD has increasingly emphasized important roles for myeloid lineage biology, with peripheral monocytes and microglia now recognized as critical modulators of neuropathology^16–18^. For these reasons, the human monocytic cell line THP-1 was used to screen for enhancer- and allele-specific transcriptional regulatory output.

In this study, we utilized an unbiased MPRA in THP-1 cells to systematically evaluate transcriptional outputs, successfully identifying 62 functional variants across 18 of the 24 (75%) tested LOAD GWAS loci. Detailed functional follow-up nominated rs636317 as the likely causal variant driving the LOAD risk association at the *MS4A* locus.

## Materials and Methods

### Identification of SNPs and design of oligos

We downloaded the summary statistics of the International Genomics of Alzheimer’s Project GWAS meta-analysis ^2^(https://www.niagads.org/igap-rv95 summary-stats-kunkle-p-value-data) and, excluding the *APOE*, *TREM2*, and *HLA* loci, selected the index SNPs from the remaining genome-wide significant loci (*P*<5e-8) for screening in the MPRA. We used the web-based tool LDlink ^19^ to identify all variants in linkage disequilibrium (r2 > 0.5) with the AD GWAS index SNPs, based on the European population data from 1000 Genomes Phase 3. All insertions, deletions, and non-bi-allelic variants were filtered out, resulting in a list of 2,231 SNPs. For each SNP, we extracted 173 bp of genomic sequence centered around the SNP using the ‘BSgenome.Hsapiens.UCSC.hg19’ annotation package in R. To each 173 bp sequence, we appended Clone1 (5’-ACTGGCCGCTTCACTG-3’) to the 5’ end and Clone2_BC (5’-GGTACCTCTAGANNNNNNNNNNNNAGATCGGAAGAGCGTCG-3’) to the 3’ end, where NNNNNNNNNNNN corresponds to a unique 12 nt barcode, and the remaining sequences are used for cloning into pMPRA series plasmid vectors. Each reference and alternative allele of the 2,231 SNPs was assigned 20 of the unique 12 nt barcodes and the resulting 89,240 designed sequences were synthesized on an Agilent Technologies 100 K 230-mer array.

### Library generation

The complex oligonucleotide pool produced from the array synthesis was amplified by emulsion PCR and cloned into the pMPRA1 (Addgene ID 49349) plasmid vector according to the established protocol ^20^. Briefly, we amplified our sequences in emulsion PCR reactions, using the MPRA_SfiI_F (5’- GCTAAGGGCCTAACTGGCCGCTTCACTG-3’) and MPRA_SfiI_R (5’- GTTTAAGGCCTCCGTGGCCGACGCTCTTC-3’) primers to add *SfiI* sites for cloning into the pMPRA1 backbone. Emulsion PCR was carried out in 350 uL reactions, with 50 uL of PCR reaction mixture containing Herculase II Fusion DNA polymerase (Agilent Technologies), and 300 uL of oil and surfactant mixture from the Micellula DNA Emulsion and Purification kit (CHIMERx) for 20 cycles. The emPCR product was cloned into an *Sfi1*- linearized pMPRA1 vector backbone with T4 DNA ligase (NEB). The ligated library was transformed into Stellar Competent cells (Takara Bio) in 8 parallel transformations. Recovered cells were pooled and used to inoculate 200 ml of LB supplemented with 100 ug/ml carbenicillin, which was placed at 37 °C and grown overnight. The amplified complex plasmid library was purified by Hi-Speed Midiprep (Qiagen) and digested with *KpnI* and *XbaI* (NEB) so that the SV40 promoter and inert *luc2* ORF donor fragment from the pMPRAdonor3 vector (Addgene ID 49354) could be inserted between the SNP-containing genomic sequence and the 12 nt barcode. The digested plasmid library and pMPRAdonor3 fragment were then ligated with T4 DNA ligase (NEB) and the resulting ligation product was transformed into competent cells, grown overnight, and purified by midiprep as described above to produce the finished MPRA plasmid library.

### Cell culture

THP-1 cells (ATCC) used for MPRA and luciferase assays were cultured in RPMI 1640 media (Corning) with 10% FBS (GeminiBio) and 1x Pen/Strep (Corning). Human ESCs (H9) were cultured in mTeSR with 5x supplement (StemCell Technology).

### MPRA screen

We transfected 2 million THP-1 cells (ATCC) with 500 ng of our library in triplicate using the Amaxa Nucleofector Kit V (Lonza), with the Y-001 program on a Nucleofector II electroporation system. Twenty-four hours post-transfection, total RNA was harvested, poly-A mRNA was isolated with magnetic beads (Thermo Fisher), cDNA was synthesized by RT-PCR, and Illumina sequencing libraries were made from both the synthesized cDNA, and aliquots of the input MPRA plasmid DNA library, specifically amplifying the unique barcode region by using the custom primers TAGseq_P1 (5’-AATGATACGGCGACCACCGAGATCTACACTCTTTCCCTACACGACGCTCTTCCGAT CT-3’) and TAGseq_P2 (5’- CAAGCAGAAGACGGCATACGAGAT[index]GTGACTGGAGTTCAGACGTGTGCTCTTCCGATCTCGAGGTGCCTAAAGG-3’), as previously described^20^. The resulting libraries were sequenced on an Illumina NextSeq 500 as a High-Output 50bp SR run by the GCB Sequencing and Genomic Technologies Core at Duke University.

### Analysis of MPRA screen

All sequencing reads were aligned to a reference FASTA file that matched each barcode to its corresponding sequence of origin, using BWA mem (Li, 2013). All low-quality barcode reads were filtered by removing any read that contained one or more position with a Phred score less than 30, or that did not exactly match a designed barcode. To determine sequence activity, the sum of the barcode counts for each designed oligo within each replicate was normalized, and oligos were tested for differential expression relative to the plasmid input through the use of a negative binomial model in DESeq2^21^. Oligos passing an FDR threshold of 0.05 were considered to be differentially expressed, or “active”. For these active sequences, we used a t test on the log-transformed RNA/plasmid ratios for each experimental replicate to determine allele specific activity, using an FDR threshold of 0.05.

### Luciferase assays

To generate the constructs for use in the luciferase reporter assays, the sequences to be tested were ordered as 200 bp oligo fragments (170 bp of genomic sequence centered around the AD GWAS SNP, to match the MPRA-tested constructs, with 15 bp of sequence at each end that overlaps with the luciferase reporter vector) from IDT technologies and cloned into the pGL4.23[*luc2*/minP] luciferase reporter plasmid(Promega) using the In- Fusion cloning system (Clontech). The firefly luciferase reporter constructs carrying SNPs to be tested were co-transfected with the Renilla luciferase pRL-CMV vector (Promega) in 1-2 million THP-1 cells at a ratio of 2 ug:40 ug Firefly:Renilla, using the Amaxa Nucleofector Kit V (Lonza), with the Y-001 program on a Nucleofector II electroporation system. Twenty-four hours after transfection Firefly and Renilla luciferase activity was measured using the Dual-Luciferase Reporter Assay system (Promega) according to the manufacturer’s protocol. Luminescence was measured by Synergy 4 multimode luminometer (Biotek) and Firefly luciferase activity was normalized to the activity of Renilla luciferase. Six technical and 3 biological repeats were performed per construct. Significance was calculated by Wilcoxon matched-pairs signed rank test.

### Tri-HiC data

Our recently developed ultra-high resolution Hi-C methodology, Tri-Hi-C, was performed on primary CD14+ monocytes of 4 Type I Diabetes patients, as part of an unpublished and unrelated study, according to our established protocol^22^. Using the BAM files from each Tri-Hi-C sample, SNP and INDEL discovery and genotyping was performed across all 4 samples simultaneously, using variant quality score recalibration, and according to GATK Best Practices recommendations^23,24^. This resulted in ∼3 million called SNPs per sample. The HiCCUPS algorithm of the Juicer Hi-C analysis toolset^25^ was used to call significant chromatin interaction loops at 10kb resolution from the contact matrices of each sample, with the parameters: -k KR -r 10000 -f 0.1 -t 0.02,1.5,1.75,2.

### Enrichment analysis

ATAC-seq data of THP-1^26^ was downloaded from the Gene Expression Omnibus (GSE96800) and ATAC-seq data of ex-vivo human microglia was downloaded using the Table Browser function of the UCSC Genome Browser (group=”Microglia PMID 28546318”, table=”hub_166047_exvivo_atac_seq_peaks”) and translated to hg19 coordinates using LiftOver. ATAC-seq peaks from these two datasets were assessed for significant overlap with “active” MPRA sequences by the Fisher’s exact test function in BEDTools^27^. Significance threshold for overlap was set as *P* < 0.05. HOMER^28^ motif enrichment analysis was performed on active MPRA sequences using findMotifsGenome.pl with default parameters. FDR significance threshold was set as *P* < 0.05.

### Variant annotation and motif disruption prediction

Using the software package BEDTools^27^, we intersected all variants found in sequences that exhibited significant allelic skew in MPRA with all transcription factor TF ChIP-seq data and DNase I hypersensitivity data available in the ENCODE project database (https://encodeproject.org), as well as enhancer (H3K27ac) and promoter(H3K4me3) histone mark ChIP-seq and ATAC-seq data from primary human microglia^14^. Monocyte macrophage eQTL data from the BLUEPRINT project^29^, the CEDAR cohort^30^, Fairfax et al. 2014^31^, Nedelec et al. 2016^32^, Quach et al. 2016^33^, Alasoo et al. 2018^34^, and Schmiedel et al. 2018^35^ were downloaded from the EMBL-EBI eQTL Catalogue(https://ebi.ac.uk/eqtl)^36^ and were queried for significant associations with MPRA allelic skew variants. Multiple testing correction was performed using the Bonferroni method, with the significance threshold for eQTL *P*-values depending on the number of genes or probes tested in each study. We used the R package motifbreakR^37^ to predict the impact of MPRA-identified functional variants on transcription factor binding, using the HOCOMOCO database of transcription factor motifs and employing a significance threshold of *P* < 5e-5.

### Allelic imbalance analysis

Aligned BAM files from CTCF ChIP-seq of CTCF in THP-1^26^ were downloaded from the NCBI Sequence Read Archive (SRP102154). FASTQ files were 101 aligned to the hg19 reference genome using Bowtie2^38^ and the resulting SAM files were analyzed for significant allelic skew by Allele-specific Binding analysis tool (ABC)^39^, with default parameters.

### Construction of gRNA Plasmid for CRISPR/Cas9 editing

The gRNA plasmid was constructed following the protocol outlined in a previous study. Briefly, three gRNA sequences were selected within the region of the rs636317 locus where Cas9 binding and cleavage occur, based on their high specificity and low off-target potential. The two primers for gRNA construction were designed as follows: the forward primer, rs636317 Insert_F: TTTCTTGGCTTTATATATCTTGTGGAAAGGACGAAACACC GACACTTTGCTGCCATCTGC, and the reverse primer, rs636317 Insert_R: GACTAGCCTTATTTTAACTTGCTATTTCTAGCTCTAAAAC GCAGATGGCAGCAAAGTGTC, which corresponds to the reverse complement of the gRNA sequence. The donor vector was constructed as follows: TACATAGAAAAACCCAACCGTCCTCAGGACACTTTGCTGCTATCTGCTGGGAAATTGTGAAAAGTACACAAAAAGTGGTTGGAG. For the PCR amplification, High-Fidelity DNA Polymerase (Phusion®) was used. The reaction conditions were as follows: initial denaturation at 98°C for 10 minutes, followed by a temperature decrease of 0.1°C/s from 95°C to 25°C, with a 1-minute hold at each 10°C decrement. The final elongation step was performed at 72°C for 10 minutes, and the reaction was held at 16°C indefinitely. After purifying the PCR products, the gRNA sequence was ligated into the gRNA vector using a 50°C, 15-minute ligation reaction (5x in-Phusion). For the transfection, the gRNA construct was introduced into ESCs in 6-well plates (cells <100 μm) using Lipofectamine Stem Cell Reagent. After colonies reached a size of >300 μm, individual colonies were picked, expanded, and sequenced to confirm the correct gRNA sequence and select the appropriate clone.

### Differentiation of Monocytes

Monocyte differentiation from hESCs was performed using the STEMdiff™ Monocyte Kit (STEMCELL Technologies). The H9 and H9-T46 cell lines were cultured according to the manufacturer’s instructions for a total duration of 16 days. Briefly, cells were first cultured in hematopoietic medium for 6 days to promote the development of monocyte lineage precursors. On day 7, the cells were switched to monocyte differentiation medium. The differentiation protocol included medium changes every 2 days. On day 0 of differentiation, following the initial 7 days of pre-differentiation, cells were collected for the mid-stage of differentiation. Additional samples were collected after 5 days for the late stage of differentiation. Following the initial 14 days of differentiation, cells were further cultured with medium changes every 2–3 days for the remaining duration. Differentiation was monitored using microscopy to observe changes in cellular morphology. Typical monocyte-like features, including increased cell size and a round shape, became evident over time. To confirm successful differentiation, the expression of monocyte-specific markers was assessed by fluorescence and qPCR. Specifically, the expression of CD14 and CD16, established markers for monocytes, was analyzed. Cells were stained with antibodies against CD14 and analyzed by fluorescence using Alex594. The differentiated monocytes from both the H9 and H9-T46 lines were confirmed to express these markers.

### Detection of MS4A Family Gene and TREM2 Expression

After the differentiation of monocytes, approximately 1 x 10^6 cells were collected by centrifugation and washed twice with PBS to remove any residual media or contaminants. Following the wash steps, cells were lysed using a lysis buffer and extrated RNA using kit (Qiagen RNeasy Kit). RNA integrity and quantity were assessed using a Nanodrop, ensuring high-quality RNA for downstream applications. For the detection of MS4A family gene expression, complementary DNA (cDNA) synthesis was performed using a reverse transcription kit (Thermo Fisher, SuperScript III) following the manufacturer’s instructions.

The expression levels of MS4A family genes and TREM2, including MS4A6E, MS4A4A, MS4A4E, MS4A6A, MS4A7, and MS4A14, were measured by quantitative PCR (qPCR). For this, primers specific to each gene were designed and optimized for qPCR conditions. PCR was performed on a Real-Time PCR System (Applied Biosystems QuantStudio 3) using the following cycling conditions: initial denaturation at 95°C for 10 minutes, followed by 40 cycles of denaturation at 95°C for 15 seconds, annealing at the appropriate temperature (typically 60°C) for 30 seconds, and extension at 72°C for 30 seconds. Melting curve analysis was performed to confirm the specificity of the PCR products. The expression of target genes was quantified relative to an internal housekeeping gene, β-actin, using the 2^(-ΔΔCt) method. The fold change in gene expression was calculated for each of the MS4A family genes of interest, comparing unedited monocytes to baseline control. Statistical significance of the differences in expression levels was determined using appropriate statistical ANOVA.

### ELISA TREM2

The TREM2 ELISA (Invitrogen) was performed to quantify human TREM2 levels cell culture supernatants. Samples were prepared by concentrating in 10x using Amicon Ultra-2ml (Milie pore). The pre-coated microplate was incubated with standards and samples, followed by detection using a biotinylated antibody and streptavidin-HRP conjugate. After substrate addition, colorimetric reaction was measured at 450 nm using a microplate reader. TREM2 concentrations in the samples were determined by comparing the absorbance values to a standard curve. The assay was conducted following the manufacturer’s instructions, with results analyzed using standard data processing software.

## Results

### Designing an MPRA to screen LOAD GWAS variants

Using the summary statistics of the recent International Genomics of Alzheimer’s Project (IGAP) LOAD GWAS meta-analysis^2^, we selected the index SNPs from the 22 genome-wide significant loci (*P*-value < 5e-8), not including the *APOE*, *TREM2*, and *HLA* loci, as well as the index SNPs from two loci (*MEF2C* and *NME8*) that had been found as genome-wide significant in the previous IGAP meta-analysis, but which did not meet the cutoff for the current study. We then used the web-based tool LDlink to select all bi-allelic SNPs in linkage disequilibrium (LD, r2 >0.5) with the LOAD GWAS SNPs, resulting in a total of 2,231 SNPs (**Figure 1a**). Oligonucleotides carrying either the reference or alternative alleles of our selected SNPs, centered in 173 bp of their genomic sequence context, were synthesized by microarray, with both alleles of every SNP assigned 20 unique 12 nucleotide barcodes (**Figure 1a**). The synthesized oligos were then cloned into pMPRA plasmid reporter vectors (**Figure 1b**). An SV40 promoter and inert *luc2* open reading frame (ORF) were inserted between the SNP carrying sequence to be tested and its assigned unique barcode (**Figure 1c**). This orientation allows for the assessment of the transcription-driving activity of the candidate SNP-harboring sequence, while incorporating the unique barcode into the 3’-UTR of the inert ORF for subsequent sequencing-based detection of activity. To measure the relative activity of each genomic sequence and its corresponding SNP, the plasmid library was transfected into the monocytic THP-1 cell line in triplicate, mRNA was isolated, and sequencing libraries were created from both the reverse-transcribed mRNA and the plasmid DNA input (**Figure 1d**). After next-generation sequencing of the barcode libraries (**Figure 1e**), the normalized log2 proportion of mRNA reads/DNA reads for each barcode-SNP allele combination was used to determine significant allelic skew in transcriptional activity (**Figure 1f**).

**Figure 1.**
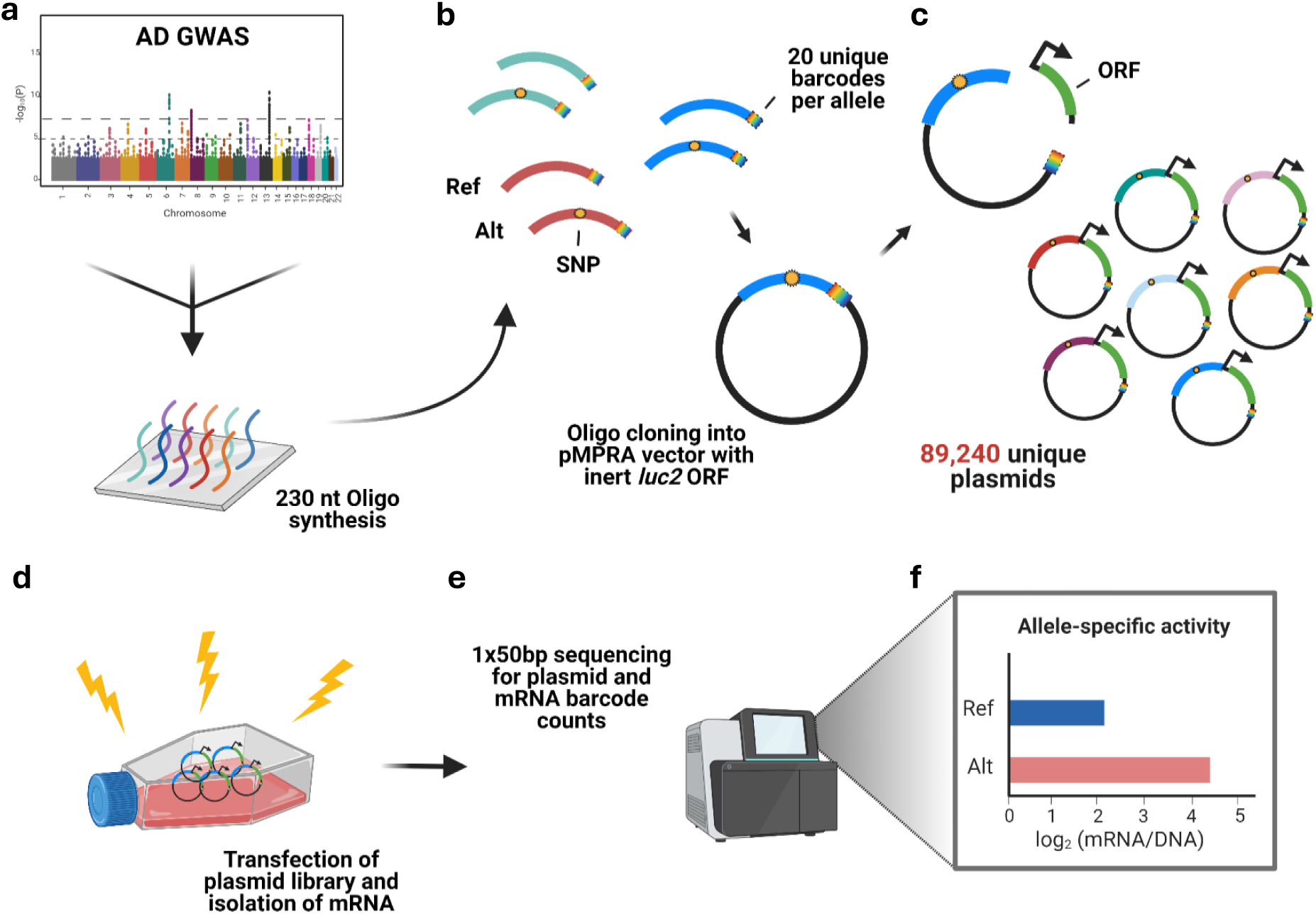
Workflow of Massively Parallel Reporter Assay. (**a**) For each AD GWAS index SNP we identified all SNPs in LD (r2 > 0.5), extracted the 173 bp of genomic sequence surrounding each SNP, and synthesized oligonucleotides containing the genomic sequence, centered on the SNP. Synthesized oligos were 230 nt in length and also contained primer adaptors and restriction sites for cloning, as well as unique 12 nt barcodes, 20 per reference and alternative allele of each SNP. (**b**) The single-stranded oligos are amplified and converted to double-stranded DNA by emulsion PCR, which is cloned into the pMPRA1 vector backbone. (**c**) The plasmid library is then linearized between the SNP-containing oligo sequence and the unique barcode by restriction digest, and a fragment from the pMPRAdonor3 vector, containing an SV40 promoter and an inert *luc2* open reading frame (ORF), is ligated in to create the final MPRA plasmid library. (**d**) The complex MPRA plasmid library is transfected into THP-1 cells by electroporation, RNA is harvested 24 hrs later from the transfected cells, and mRNA is isolated. (**e**) Illumina libraries are prepared from both harvested mRNA and plasmid library input and sequenced as a 1x50bp run on a NextSeq 500. (**f**) Sequenced barcode counts are matched to their respective oligo/SNP construct and allele-specificity of reporter activity for each SNP is determined by comparing the log-transformed ratio of mRNA/plasmid DNA barcode counts.

### Determining regulatory activity of the oligos

Before identifying potential allelic skew, we first identified the subset of genomic sequences for which at least one of the two alleles showed significant transcriptional activity. We compared the expression of each allelic version of each genomic sequence to the distribution of expression of every other sequence and calculated relative activity scores for all the tested sequences. Out of 4,462 tested sequences, 975 (22%) were determined to be “active” (**Figure 2a**). Since MPRA tests the activity of genomic elements outside of their native chromatin environment, we examined the overlap of our active sequences with regulatory elements identified in THP-1 cells^26^ and primary microglia^40^. We found that these active sequences were significantly enriched for open chromatin, as measured by ATAC-seq analysis, compared to the sequences determined to be inactive, or to 10,000 size- and GC content-matched random genomic sequences (**Figure 2b**). When we performed motif enrichment analysis of the active sequences we identified enrichment of transcription factor (TF)-binding motifs known to be involved in myeloid biology, including THRa/THRb^41^, LXR^42^, and RARa^43^ (**Figure 2c**). Together, this provided evidence that MPRA can identify regulatory sequences that are active in the native chromatin setting and suggested that differential MPRA activity could represent alterations in the binding or activity of TFs active in the myeloid lineage.

**Figure 2.**
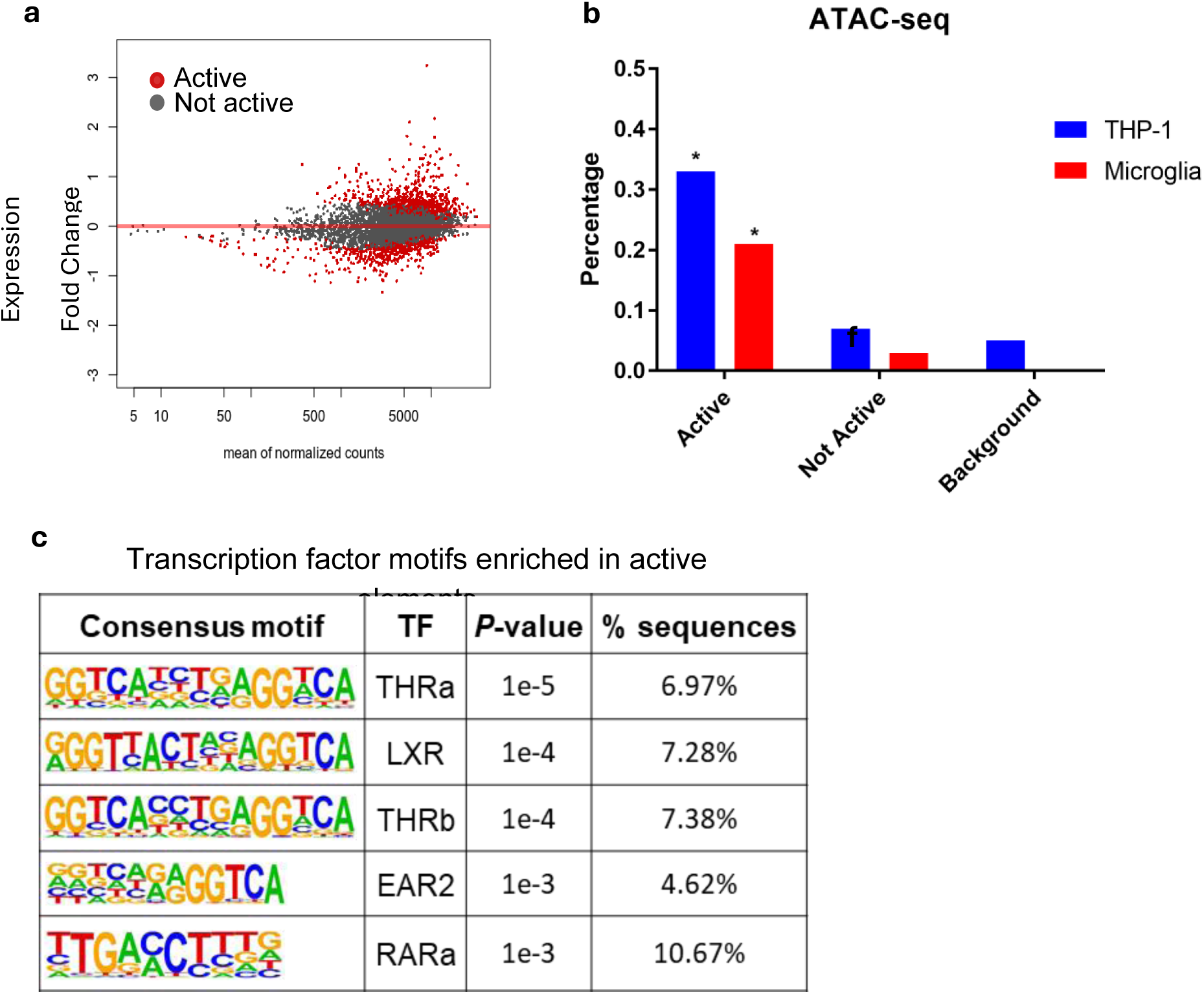
MPRA in THP-1 identifies active regulatory elements that are enriched for open chromatin and myeloid TF motifs. (**a**) 975 of 4,462 (22%) sequence tested by MPRA were found to be active by DESeq2. Red dots correspond to active sequences and are plotted as expression fold change log2 (RNA/Plasmid) vs mean of normalized sequence counts (**b**) MPRA active sequences are enriched in ATAC-seq peaks in THP-1 and primary microglia, compared to non-active sequences and 10,000 random genomic background sequences (**c**) MPRA active sequences are enriched for myeloid TF motifs. \**P* < 0.05.

### Allele-specific regulatory activity of LOAD SNPs in MPRA

Taking into consideration only those sequences which we had determined to be active, we next set out to identify if any exhibited significant allelic variation in activity. Out of the 975 active sequences, 62 showed significant allelic skew (**Figure 3a**), representing 18 out 24 (75%) LOAD GWAS loci tested. To determine potential mechanisms underlying this allelic skew we investigated the overlap between the variants corresponding to each sequence and TF ChIP-seq and DNase I hypersensitivity data from all available cell types in ENCODE^44^, as well as enhancer (H3K27ac) and promoter (H3K4me3) ChIP-seq and ATAC-seq data from primary microglia^14^, and predicted the variants’ ability to disrupt TF motifs, with the assumption that allelic skew in sequence activity is mediated by effects on TF binding to regulatory elements. To identify potential target genes, we also queried monocyte and macrophage expression quantitative trait loci (eQTL) data from several studies^29–35^ for associations with MPRA allelic skew variants. As a result, we found strong functional evidence for 25 of the 62 variants. We then validated the allele-specific transcriptional effects of three of the variants supported by strong functional evidence by traditional luciferase reporter assay (**Figure 3c**), with all results concordant with the MPRA findings. Increased reporter expression was observed from the risk allele of rs636317 (**Figure 3b-c**), which resides in an intergenic region in the *MS4A* risk locus marked by open chromatin, disrupts a binding motif for the structural chromatin factor CTCF (**Figure 3d**), and is also a highly significant eQTL for *MS4A6A*, *MS4A4A*, *MS4A4E*, and *MS4A6E* in monocytes (**Table 1**).

**Figure 3.**
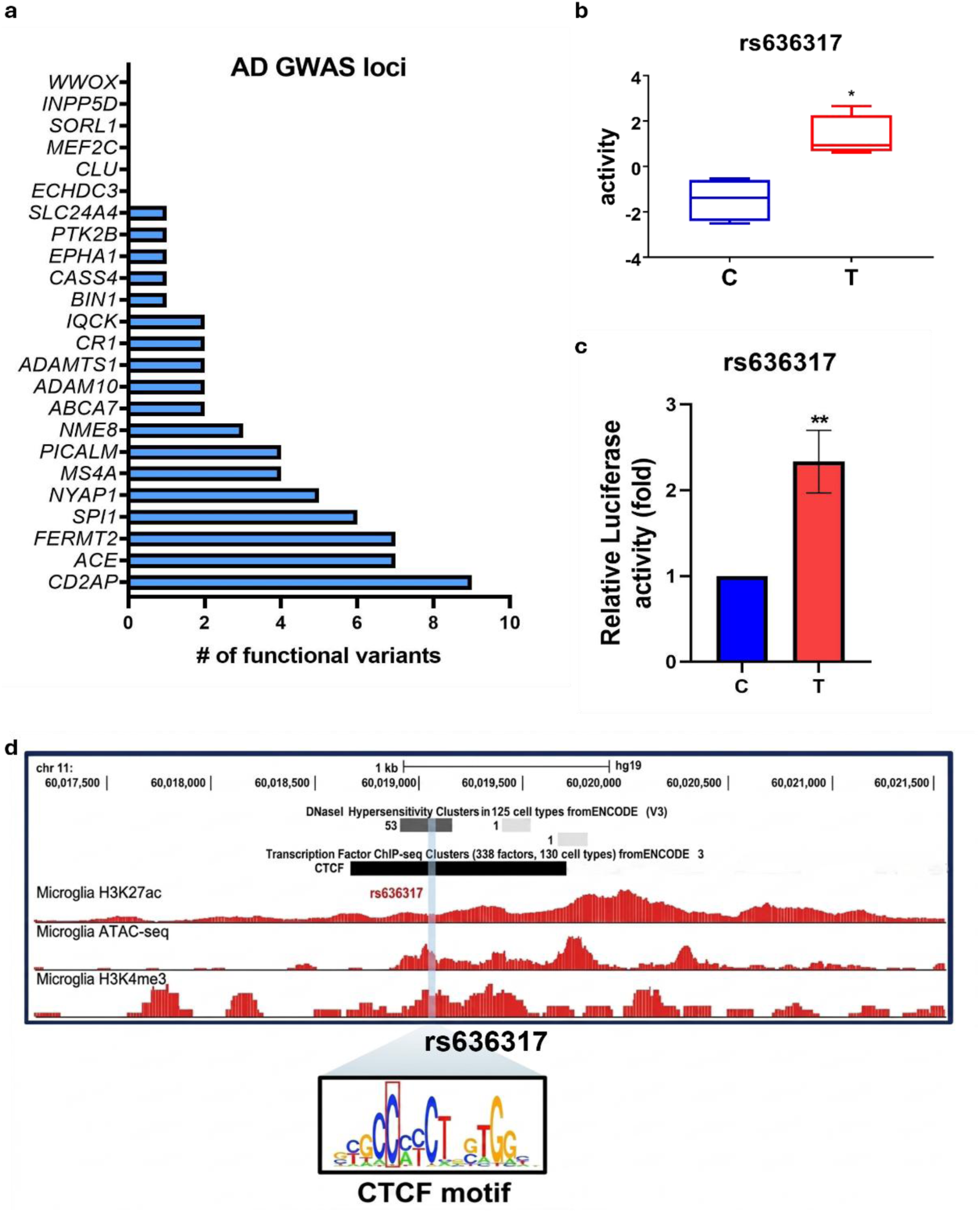
MPRA in THP-1 identifies allele-specific effects of LOAD GWAS SNPs. (**a**) Functional variants demonstrating allelic skew in activity are found for 18 of 24 LOAD GWAS loci tested by MPRA (**b**) MPRA identifies significant allelic skew in activity for the *MS4A* locus SNP rs636317 (**c**) rs636317 allelic skew is validated by luciferase reporter assay in THP-1 (**d**) rs636317 resides in an open chromatin region in 53 ENCODE cell types, is bound by CTCF in 137 ENCODE cell types and also resides in a region of open chromatin marked by H3K27ac in primary human microglia. \**P* < 0.05, \*\**P* < 0.01.

**Table 1.** rs636317 eQTL genes in monocytes.

| Gene | Probe ID | Study | Monocyte |  |
| --- | --- | --- | --- | --- |
|  |  |  | P value | Beta |
| *MS4A6A* | — | BLUEPRINT | 1.38E-09 | 0.089079 |
| *MS4A4A* | — | BLUEPRINT | 1.73E-15 | 0.612692 |
| *MS4A4E* | — | BLUEPRINT | 1.82E-06 | 0.181486 |
| *MS4A6A* | ILMN_1721035 | CEDAR | 9.48E-11 | 0.126126 |
| *MS4A6A* | ILMN_2359800 | CEDAR | 1.39E-07 | 0.099815 |
| *MS4A4A* | ILMN_2370336 | CEDAR | 6.60E-07 | 0.141246 |
| *MS4A6E* | ILMN_1759933 | CEDAR | 9.62E-06 | 0.041628 |
| *MS4A6A* | ILMN_1721035 | Fairfax 2014 | 3.69E-13 | 0.063077 |
| *MS4A4A* | ILMN_2370336 | Fairfax 2014 | 1.50E-12 | 0.157429 |
| *MS4A4A* | — | Schmiedel 2018 | 2.87E-06 | 0.283037 |

We chose to investigate the mechanism of action at rs636317 in more detail due to the fact that *MS4A* locus genes have been implicated in LOAD pathological processes involving *TREM2*^45^, a well-recognized LOAD risk gene with important roles in myeloid cell biology^46^, as well as the fact that alterations to CTCF sites have been implicated in multiple diseases and conditions^47,48^. Examination of DNase I hypersensitivity and TF ChIP-seq data from ENCODE found that rs636317 resides in a region of open chromatin with conserved CTCF binding in many, if not all, cell types (**Figure 4a**). Several other TFs also show binding to the region, including the cohesin complex components SMC3 and RAD21, and the zinc finger TF ZNF143, all of which play key roles in the formation of promoter-enhancer chromatin loops^49^. Since rs636317-T was predicted to disrupt CTCF binding, we wondered whether genotype-dependent effects could be seen in CTCF binding in cell types assayed for CTCF ChIP-seq in the ENCODE data. As rs636317 was not genotyped by ENCODE, we used the highly linked SNP rs1562990 as a proxy for rs636317 genotype (*r2* =0.99, D’=1, EUR population 1KG). Examination of CTCF peaks in CD14+ Monocytes and GM12878 lymphoblastoid cells, both homozygous for rs636317-C, found a robust peak at the motif, and CTCF ChIA-PET data from GM12878 showed an anchoring of upstream chromatin interactions at rs636317, with no interactions observed with elements downstream (**Figure 4b**), indicating that the CTCF binding site at rs636317 might act as a topologically associating domain (TAD) boundary element. Hi-C data from GM12878 and IMR90 also supported a TAD boundary role for rs636317. Another rs636317-C homozygous cell line, K562, also showed a robust peak at the motif, while the rs636317-T homozygous IMR90, HRE, and NHDF cell types exhibited a lack of CTCF binding (**Figure 4b**). Evidence of CTCF binding in two cell types with the heterozygous rs636317-C/T genotype was mixed, with NH-A cells showing no binding, and SK-N-SH cell line showing a weak peak at the motif (**Figure 4b**). While these observations provided qualitative evidence of genotypic effects on CTCF binding at rs636317, we next sought to gain quantitative evidence. By performing sequencing read-based allele109 specific binding analysis, using CTCF ChIP-seq data from the heterozygous SK-N-SH cell line, we observed a clear and highly significant allelic skew in CTCF binding at rs636317 (**Figure 4c**). This result is supported by recently published work which also nominated rs636317 as a candidate causal variant in the *MS4A* locus, and similarly found significant allelic skew in open chromatin, as measured by ATAC-seq in iPSCderived microglia^15^.

**Figure 4.**
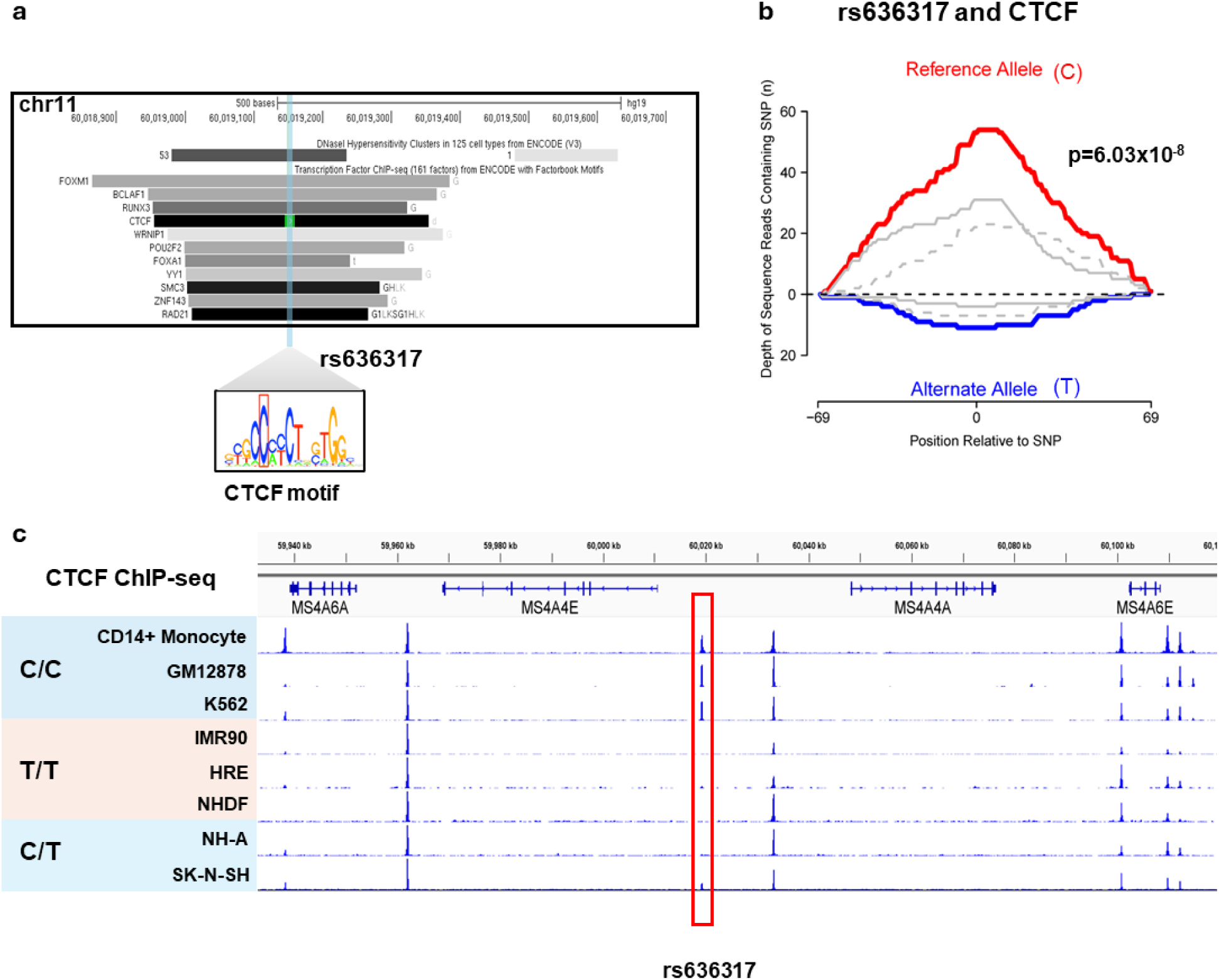
*MS4A* locus SNP rs636317 exhibits allele-specific effects on CTCF binding. (**a**) rs636317 is bound by structural chromatin proteins involved in loop formation (**b**) rs636317 shows allele-specific effects on CTCF binding in ENCODE ChIP-seq data (**c**) rs636317 exhibits significant allelic skew in CTCF binding in heterozygous SK-N-SH cells.

### Causal variant in the *MS4A* locus alters chromatin loop mediated interactions

All of the evidence we had compiled thus far indicated that rs636317 risk genotype was most likely affecting LOAD risk by impacting large-scale TAD structure in the *MS4A* locus, thus resulting in alterations to local gene expression, specifically in myeloid cells. However, allele-specific binding of CTCF to rs636317 might also elicit more subtle changes to chromatin looping in the locus, such as alterations to promoter-enhancer interactions which do not affect larger-scale structures. Using our ultra-high resolution chromatin conformation capture method, Tri-Hi-C^22^, we have recently profiled the three-dimensional chromatin interaction landscape of human primary T cells and monocytes from 4 individuals (unpublished data), with a resolution that allowed us to examine monocyte chromatin interactions in more detail than had been achieved in previous Hi-C studies^26^ (**Supplementary Figure 1**). As the publicly available chromatin conformation data was not of sufficient resolution to allow for the detection of such subtle changes to chromatin looping, we decided to re-analyze our previously generated monocyte Tri-Hi-C data to identify allele-specific effects brought about by rs636317 genotype. Since Tri-Hi-C is a genome-wide sequencing assay, we were able to perform variant calling and genotyping for all of our samples, determining that two of four individuals in our dataset were heterozygous for rs636317, while the remaining two were homozygotes for the risk allele rs636317-T which disrupts the CTCF binding site. We performed loop calling on the Hi-C matrix of each sample and found that there were significant differences in loop-mediated interactions between genotypes (**Figure 5**). Given the evidence from the eQTL data, *MS4A6A*, *MS4A4A*, *MS4A4E*, and *MS4A6E* are the genes most likely to be affected by rs636317 genotype. We did not observe a gain or loss of loop-mediated interactions with the *MS4A6A* or *MS4A4E* promoters between genotypes, but we did observe a gained interaction between the *MS4A4A* promoter and a CTCF-bound enhancer element ∼25kb upstream of *MS4A7* in one of the two rs636317-T homozygotes (**Figure 5**). We also identified gained interactions between the CTCF site at rs636317 and an enhancer element ∼100kb downstream, as well as between rs636317 and a CTCF anchor ∼350kb downstream (**Figure 5**), in the absence of the protective allele rs636317-C.

**Figure 5.**
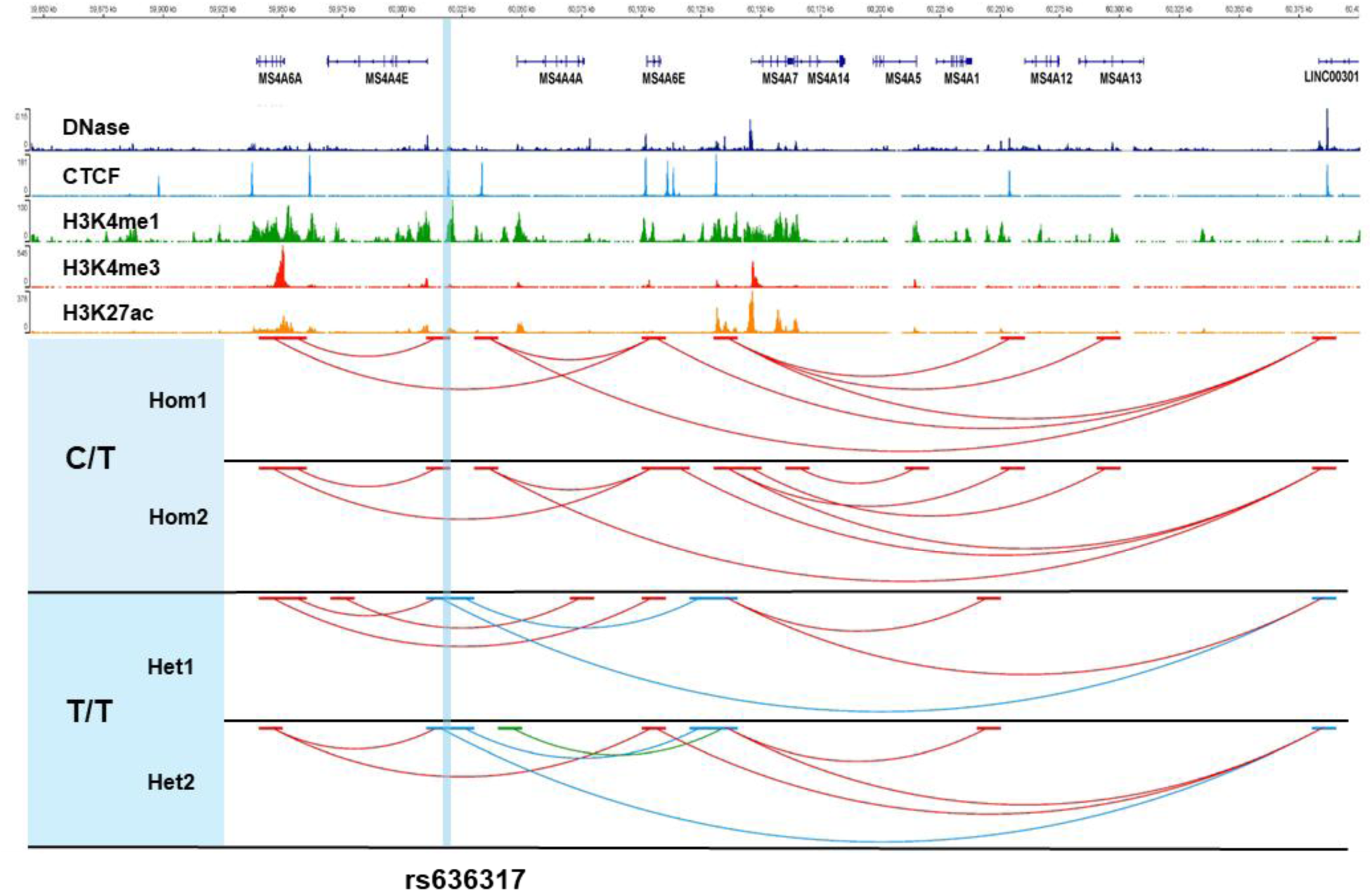
rs636317 alters chromatin loop-mediated interactions in the *MS4A* locus. (**I**) ENCODE ChIP-seq and DNase-seq tracks from CD14+ Monocytes indicate the location of open chromatin regions, CTCF binding sites, and enhancer (H3K4me1/H3K427ac) and promoter (H3K4me3/H3K27ac) regions. (**II**) Significant chromatin loops called from Tri-Hi-C data of two individuals heterozygous for rs636317 genotype (**III**) Significant chromatin loops called from Tri-Hi-C data of two individuals homozygous for rs636317-T. Blue shaded bar indicates the location of the CTCF binding site occupied by rs636317. Yellow shaded bars indicate the promoter of *MS4A4A* and a CTCF-bound enhancer region upstream of *MS4A7*. Chromatin loop gain at rs636317 can be seen in both heterozygous individuals (**III**), and gain of interaction at the *MS4A4A* promoter can be seen in the heterozygous individual labeled 1035_Mono_rs636317_TT.

### Allele-specific deletion of rs636317 enhances expression of MS4A4E and MS4A6A

To directly validate the predicted effects of the rs636317 genotype on gene expression in the MS4A locus, we generated a monoclonal-edited hESC model. Using CRISPR/Cas9 technology, we specifically targeted the rs636317 region in H9 hESCs to create a heterozygous deletion of 10 nucleotides within the CTCF binding motif associated with the rs636317-T risk allele (**Figure 6a**). The resulting cell line, designated T46, was confirmed by sequencing, and off-target effects were minimized **(Supplementary Table 1**). Consistent with our MPRA and eQTL findings, we differentiated the T46 hESCs into monocytes to assess the transcriptional impact of the rs636317-T deletion. Among the 18 members of the human MS4A family, we observed that monocyte-derived T46 cells exhibited significantly increased expression of MS4A4E and MS4A6A (**Figure 6b**). These results confirm that the rs636317-T allele, which disrupts CTCF binding, leads to the upregulation of specific MS4A family transcripts.

**Figure 6.**
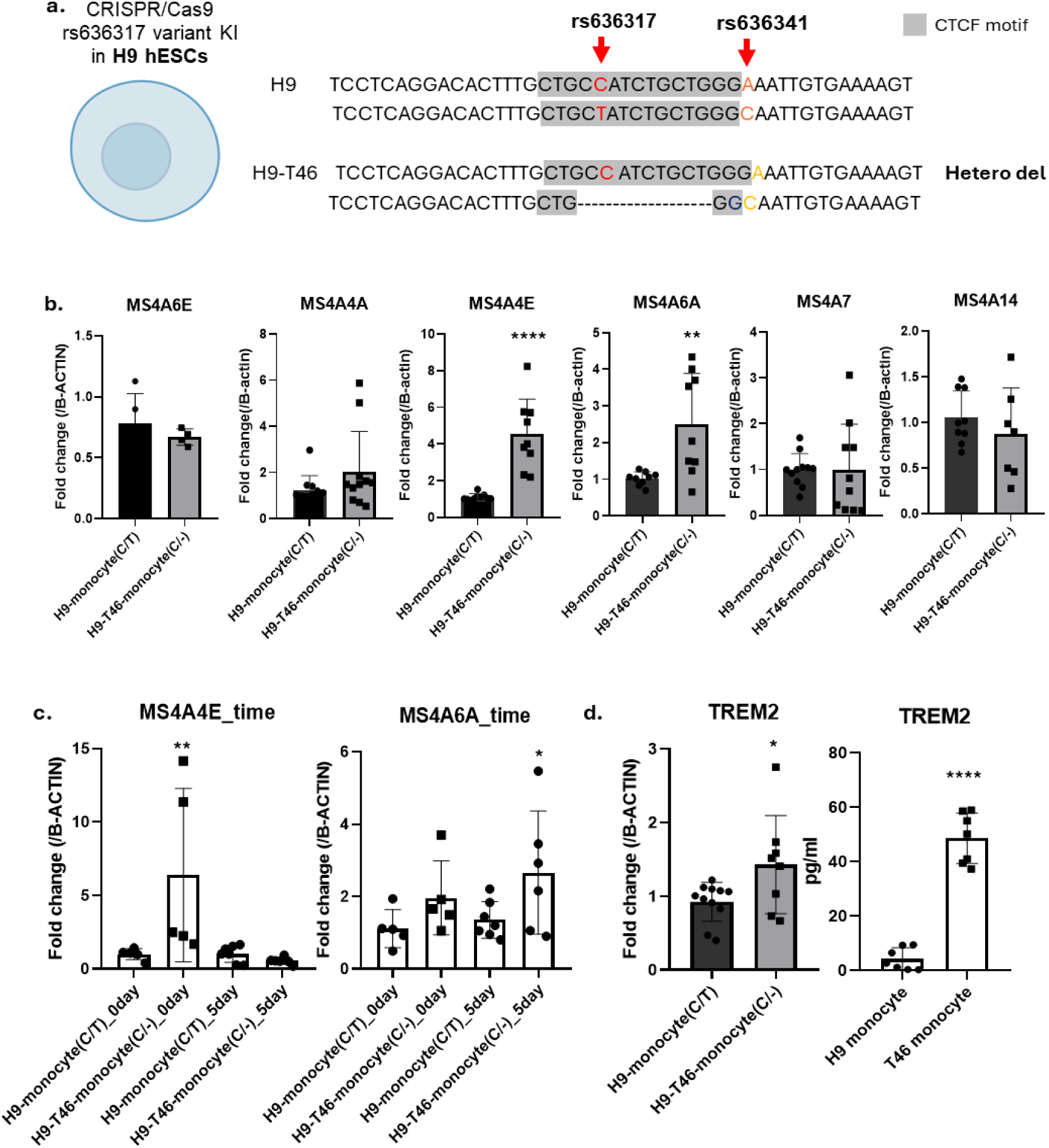
Deletion of the rs636317 T variant enhances the expression of MS4A4E, MS4A6A, and TREM2. a) Schematic representation of the variant deletion in the cell line. The rs636317 T variant was deleted in human ESC-H9 cells using CRISPR/Cas9, resulting in the generation of the heterozygous deletion line, H9-T46. b) The expression levels of MS4A family genes (including MS4A6E, MS4A4A, MS4A4E, MS4A6A, MS4A7, and MS4A14) were assessed by qPCR, revealing a significant increase in MS4A4E and MS4A6A expression. c) During monocyte differentiation, the expression of MS4A4E and MS4A6A varied over time. d) The transcript and protein levels of TREM2 were significantly elevated in the T deletion line.

We further analyzed the expression patterns of these target genes across a 16-day differentiation protocol (7 days to hematopoietic stem cells; 9 days to monocytes). MS4A4E expression increased primarily during the early stages of differentiation. MS4A6A expression was more prominent at the mid-stage of differentiation (**Figure 6c**).

### Coordination with TREM2 signaling

Previous genetic analyses have linked variants in the MS4A locus to levels of soluble TREM2 (sTREM2) in the cerebrospinal fluid^46,50^. Specifically, variants that increase MS4A4A and MS4A6A expression are known to modulate sTREM2, which serves as a biomarker for Alzheimer’s disease (AD) progression^45,50^. Following differentiation of the T46 line into monocytes, we assessed the regulation of TREM2. We observed a 1.5-fold increase in TREM2 mRNA (**Figure 6d**). Analysis of conditioned media revealed a ∼10-fold increase in sTREM2 protein levels (**Figure 6d**). These findings suggest a mechanistic link where the rs636317-T risk allele regulates MS4A4E and MS4A6A, which in turn stimulates TREM2 expression and cleavage, potentially modulating AD risk through myeloid cell-mediated proteostasis^51^.

## Discussion

Uncovering the causal variants and effector genes underlying genetic associations is critical to realizing the potential of genome-wide association studies (GWAS) and identifying therapeutic targets for late-onset Alzheimer’s disease (LOAD). In this study, we functionally evaluated the regulatory effects of 2,231 SNPs potentially underlying 24 LOAD GWAS associations. Massively Parallel Reporter Assay (MPRA) analysis of ∼175 bp genomic sequences revealed that 22% of the tested sequences significantly modulated transcription relative to the baseline activity of the plasmid library. Allele-specific assessment identified significant regulatory skew for 62 SNPs, representing 75% of the LOAD GWAS loci investigated. Notably, over half of these loci contained multiple functional variants exhibiting allelic skew, suggesting a complex regulatory landscape at these genomic regions.

### CTCF Disruption and Chromatin Reorganization at the MS4A Locus

Through traditional luciferase reporter assays, we validated allele-specific effects for three SNPs predicted to alter transcription factor (TF) binding. Specifically, the risk allele of the MS4A locus variant rs636317 demonstrated a potent disruptive effect on CTCF binding across multiple cell types. This align with recent post-GWAS analyses nominating rs636317 as a causal variant and reporting significant allelic imbalance in open chromatin within iPSC-derived microglia and bulk brain tissue, as determined by ATAC-seq^15^. Our Tri-Hi-C analysis of primary monocytes supports a model in which the reduction or loss of CTCF binding at rs636317 alters chromatin looping. This likely creates an environment of increased interaction between local regulatory elements, thereby potentiating the transcription of nearby genes. Beyond local enhancer potentiation, our data suggest the possibility of novel chromatin interactions with MS4A member promoters, although a specific looping event between a downstream regulatory element and the MS4A4A promoter was observed in only one of two rs636317 heterozygous individuals.

### MS4A4A, sTREM2, and Myeloid Pathogenesis

The 18-member MS4A gene family remains largely uncharacterized, yet emerging evidence links several members to LOAD biology^52^. MS4A4A colocalizes within plasma membrane lipid rafts with TREM2, a pivotal regulator of myeloid function^45,46^. Our findings suggest that the rs636317-T allele confers neuroprotection by enhancing the expression of MS4A4A and MS4A6A, which in turn upregulates soluble TREM2 (sTREM2) levels. While protective variants (rs636317) correlate with increased sTREM2 and reduced AD risk, the minor allele of rs636317 is associated with diminished sTREM2 and elevated risk. These data reinforce the hypothesis that genetic modulation of immune cell function significantly influences neurodegenerative trajectories. Although LOAD GWAS variants are enriched in the enhancers of microglia, monocytes, and macrophages, the specific contribution of peripheral myeloid cells remains an area of active investigation^53,54^. In transgenic AD models, circulating monocytes effectively infiltrate the brain and exhibit superior phagocytic clearance of amyloid-beta (Aβ) compared to resident microglia^55^. Conversely, monocytes from LOAD patients show impaired recruitment and deficient Aβ phagocytosis^54,56^. For instance, the CD33 risk allele—a known repressor of immune activation—upregulates CD33 expression and decreases phagocytic capacity^57,58^. Thus, while microglial biology is central to LOAD, the role of peripheral myeloid cells is a critical, non-exclusive component of pathogenesis.

### Technical Limitations

A limitation of this study is the artificial nature of testing small genomic sequences for regulatory capacity outside of their native chromatin environment. Consequently, it is possible that there were both false positive and false negative findings in our results. Nevertheless, the enrichment of highly active sequences in open chromatin regions and myeloid-specific TF motifs supports the biological relevance of our findings. However, we did find that the most highly active genomic sequences in our assay were significantly enriched for open chromatin regions found in myeloid cell types, as well as for TF binding motifs with known roles in myeloid biology. These results give us confidence in our findings, as least with respect to the sequences and variants that are active in myeloid cells. As we were agnostic about which loci would exhibit activity in THP-1 cells, it is thus unsurprising that for some loci we did not identify any functional variants. For example, in the CLU risk locus the gene CLU itself, encoding the secreted apolipoprotein clustering, is presumed to be the causal gene. CLU is expressed in the brain primarily by astrocytes, where it has known function^59^. Consistent with a lack of microglial expression, we did not detect any functional variants in the CLU locus by MPRA in THP-1.

### Challenges in Microglial Differentiation

To investigate the MS4A family in a more relevant cellular context, we attempted to differentiate T46 embryonic stem cells (ESCs) into microglia. However, we were unable to detect MS4A expression due to unsuccessful differentiation, likely caused by reduced cell viability following CRISPR/Cas9 knock-in (KI). The deletion of rs636317 within the CTCF-binding motif may have disrupted chromatin organization essential for stem cell maintenance^49,60^. Furthermore, the genomic instability often introduced by CRISPR/Cas9 may have increased susceptibility to apoptosis or spontaneous differentiation under stress. These observations suggest that regulatory elements within the MS4A locus may influence not only mature gene expression but also fundamental cellular viability and lineage potential.

## Notes

### Competing Interest Statement

The authors have declared no competing interest.

### Funding

- NIH Common Fund: NIH AG017242, DK127778, AG076040, AG069750, AG061521, GM104459, AG056278, AG057341, AG057433, and AG057706
- Simons Foundation

