## Supplementary material for "Identification of functional non-coding variants affecting Alzheimer’s disease risk by Massively Parallel Reporter Assay": SupplementaryData.pdf

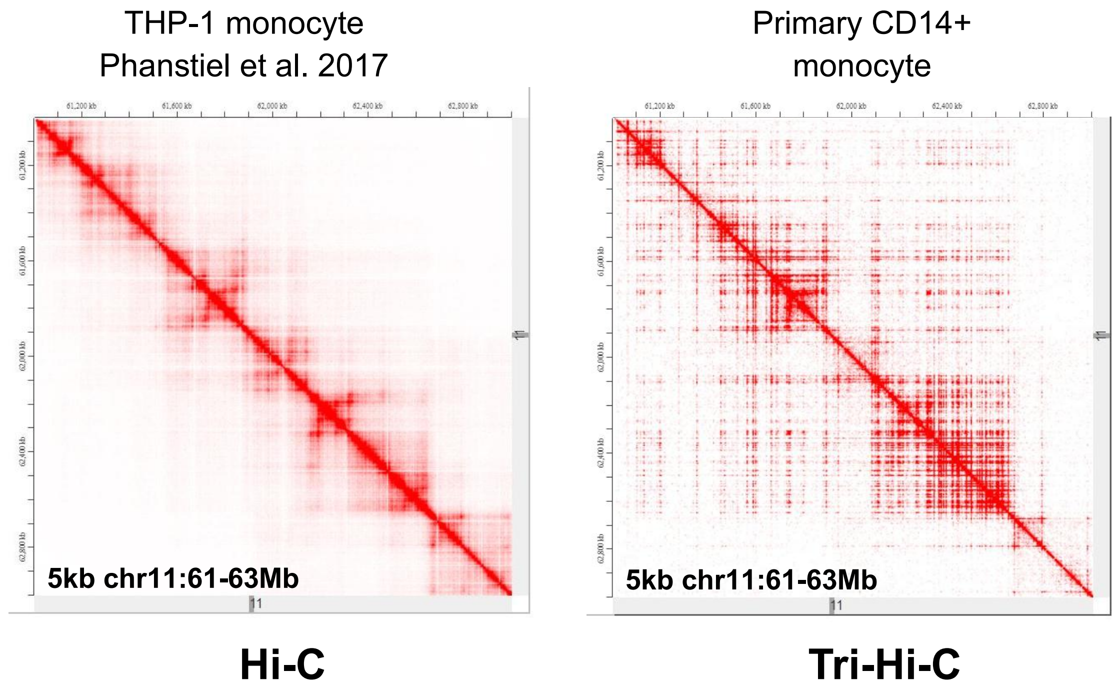


Supplementary Figure 1. Tri-HiC identifies chromatin looping with enhanced resolution

| Sequence | PAM | Mismatch | Gene | Locus | Alignment |
|---|---|---|---|---|---|
| GACACTTTGCTGCCATCTGC | CAG | | | chr6:-111652339 | Query 1 GACACTTTGCTGCCATCTGC 20<br>Sbjct 69549 GACACTTTGCTGCCATCTGC 69530 |
| GACACTTTGCTGCCATCTGC | TGG | | | chr11:-60363649 | Query 1 GACACTTTGCTGCCATCTGC 20<br>Sbjct 54 GACACTTTGCTGCCATCTGC 35 |
| GACACTTTGCTGCCATCTGC | CGG | | | chr8:+36141347 | Query 1 GACACTTTGCTGCCATCTGC 20<br>Sbjct 53810 GACACTTTGCTGCCATCTGC 53791 |
| GACACTTGGCTGCCATCTGC | TAG | 1 | | chr11:+36222727 | Query 1 GACACTTGGCTGCCATCTGC 20<br>Sbjct 17370 GACACTTGGCTGCCATCTGC 17389 |
| GGCGCTTTGCTGCCATCTGC | TGG | 2 | | chr8:+143902669 | Query 1 GGCGCTTTGCTGCCATCTGC 20<br>Sbjct 169225 GGCGCTTTGCTGCCATCTGC 169206 |
| GATA-TTTGCTGCCATCTGC | CGG | 2 | | chr9:-36798433 | Query 1 GATATTTGCTGCCATCTGC 19<br>Sbjct 16144 GATATTTGCTGCCATCTGC 16126 |
| AACCCTTGGCAGCCATCTGC | AAG | 4 | | chr3:-66825367 | Query 1 AACCCTTGGCAGCCATCTGC 20<br>Sbjct 60393 AACCCTTGGCAGCCATCTGC 60412 |

Supplementary Table 1. No off-target effect of gRNA for rs636317 T variants deletion

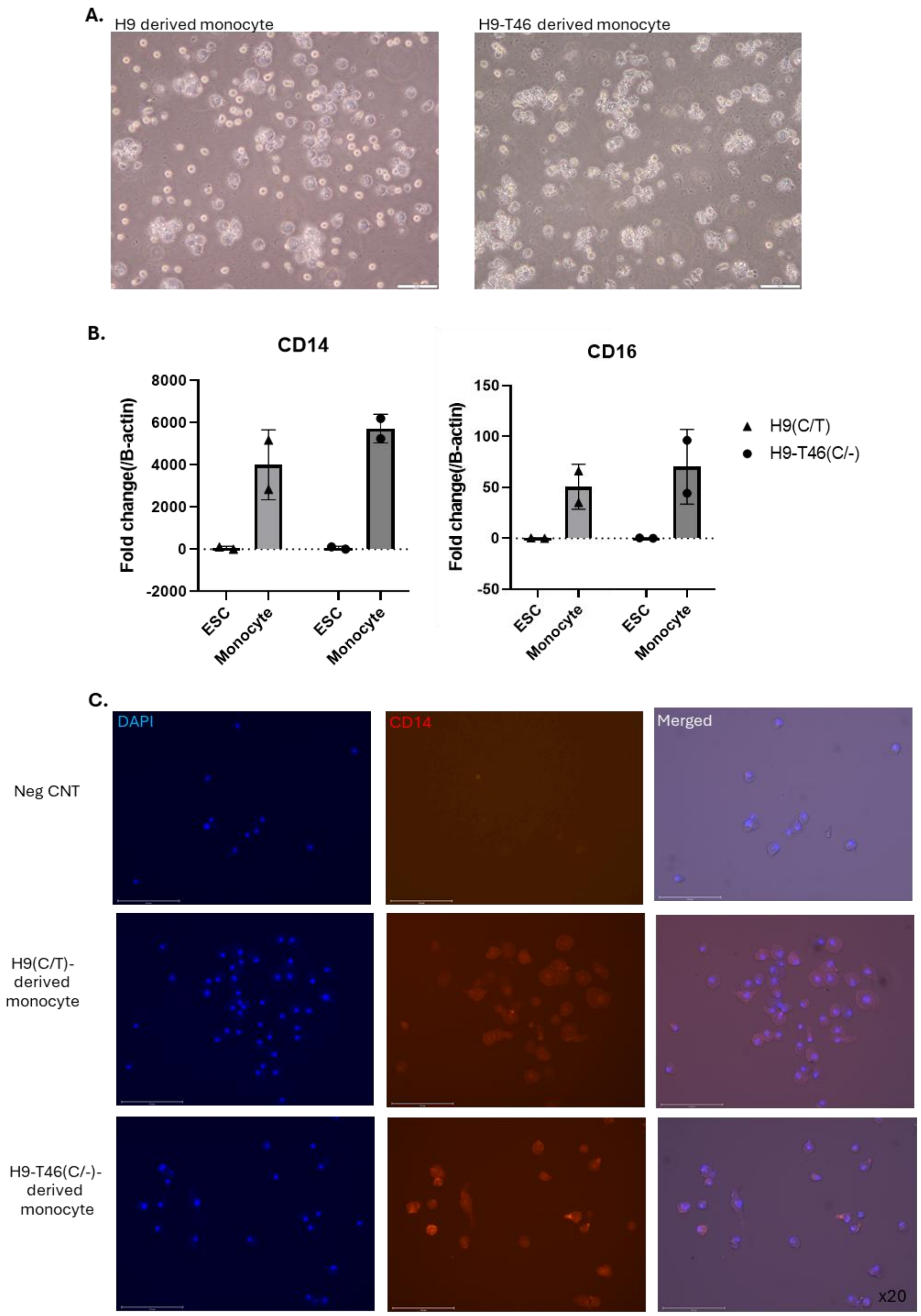


Supplementary Figure 2. Expression of monocyte markers in hESC-derived monocytes.
A) Morphological analysis confirmed the monocyte phenotype.
B) The expression of monocyte markers, CD14 and CD16, was quantified using qPCR.
C) Monocyte markers were detected in both the H9 and H9-T46 derived cell lines using CD14 antibody.
